# Ser500 phosphorylation acts as a conformational switch to prime eEF-2K for activation

**DOI:** 10.1101/2025.06.30.662482

**Authors:** Amanda L Bohanon, Luke S Browning, Rae M Sammons, Andrea Piserchio, Clint D J Tavares, Eun Jeong Cho, Ranajeet Ghose, Kevin N Dalby

**Affiliations:** Interdisciplinary Life Sciences Graduate Program, the University of Texas, Austin, TX, 78712; Targeted Therapeutic Drug Discovery and Development Program, the University of Texas, Austin, TX, 78712; Department of Chemistry and Biochemistry, the City College of New York, New York, NY 10031; Division of Chemical Biology and Medicinal Chemistry, the University of Texas, Austin, TX, 78712; PhD Program in Biochemistry, The Graduate Center of CUNY, New York, NY 10016; PhD Program in Chemistry, The Graduate Center of CUNY, New York, NY 10016; PhD Program in Physics, The Graduate Center of CUNY, New York, NY 10016

**Author notes:** A.L.B and L.S.B contributed equally to this work.

## Abstract

Eukaryotic elongation factor-2 kinase (eEF-2K), a member of the α−kinase family of atypical kinases, phosphorylates eukaryotic elongation factor 2 (eEF-2), thereby inhibiting ribosomal translocation and downregulating translational elongation in response to diverse cellular cues. eEF-2K is activated by Ca²⁺/calmodulin (CaM) and integrates upstream inputs from diverse signaling pathways, including PKA and mTOR, which target regulatory sites on a disordered regulatory loop. Among these, serine 500 (S500) has been identified as a key phosphorylation site targeted by both eEF-2K and PKA. However, the influence of this post-translational modification on the properties of eEF-2K has remained unclear. Prior studies have shown that S500 phosphorylation accelerates autophosphorylation of eEF-2K at its primary activating site, threonine 348 (T348). Here, we demonstrate that S500 phosphorylation, mimicked by a S500D mutation, works in conjunction with T348 phosphorylation to enhance the intrinsic (CaM-independent) activity of eEF-2K. Hydrogen-deuterium exchange mass spectrometry reveals that CaM binding, and consequent enhancement in eEF-2K activity, is accompanied by conformational changes proximal to S500. Deletion of S500 and surrounding residues mimics the effects of S500D, promoting robust CaM-independent activity. These data suggest that CaM binding or S500 phosphorylation have similar effects, likely relieving an inhibitory constraint to enhance activity. Further, S500 phosphorylation enhances binding to both apo-CaM and Ca^2+^/CaM, suggesting a mechanism for maintaining basal activity and priming the kinase for rapid reactivation in response to Ca^2+^ transients. These findings support a model in which phosphorylation on T348 and S500 synergize to stabilize the active conformation of eEF-2K.

## Introduction

Protein synthesis is one of the most energy-intensive processes in the cell, and its regulation is crucial for maintaining energy homeostasis and enabling adaptive responses to environmental changes (1). Cells fine-tune global translation rates and selectively control the translation of specific mRNAs to reshape the proteome in response to physiological needs (2, 3). A key regulatory node in this process is translation elongation, which, in most eukaryotes, is primarily modulated by eukaryotic elongation factor 2 kinase (eEF-2K) (4, 5). By phosphorylating elongation factor 2 (eEF-2), eEF-2K inhibits ribosomal translocation, thereby slowing elongation (6). This mechanism is thought to conserve energy, enhance translational fidelity, and prioritize the translation of stress-responsive mRNAs (7, 8). In neurons, eEF-2K plays a crucial role in synaptic plasticity, and its dysregulation has been linked to several neurological disorders (9–12).

eEF-2K is a serine/threonine kinase of the atypical α−kinase family, uniquely regulated by calmodulin (CaM) in response to intracellular Ca²⁺ signals (4, 13). Activation is initiated by Ca²⁺/CaM binding, followed by rapid autophosphorylation at T348, which stabilizes the active conformation (14, 15). Beyond Ca²⁺ signaling, eEF-2K integrates inputs from pathways such as mTOR and PKA, as well as cues related to pH, nutrient status, and cellular stress (16–22). Many of these regulatory phosphorylation sites cluster within a predicted disordered regulatory loop (R-loop), including S500, a key site phosphorylated by both eEF-2K (autophosphorylation) (15), and PKA (20, 21, 23), although its functional role remains unclear.

Here, using biochemical, biophysical, and cellular approaches, and an S500D mutant of eEF-2K, a *bona fide* mimic for S500 phosphorylation (23), we demonstrate that S500 phosphorylation synergizes with T348 autophosphorylation to significantly enhance the CaM-independent activity of eEF-2K and increase its ability to interact with both apo-CaM (Ca²⁺-free CaM) and Ca^2+^/CaM. These properties allow eEF-2K to maintain basal activity in resting cells, while remaining primed for rapid reactivation upon a Ca²⁺ influx. Our findings support a model in which S500 phosphorylation acts as a conformational switch, enabling eEF-2K to integrate Ca²⁺/CaM signals, autophosphorylation events, and upstream regulatory inputs to dynamically control translational elongation.

## Results

### S500D and T348 phosphorylation cooperatively enhance the calmodulin affinity of eEF-2K

To investigate how phosphorylation at S500 influences CaM binding, we employed a native gel electrophoresis assay using CaM labeled with the fluorophore, IAEDANS (CaM_IAE_), a construct we previously utilized to monitor eEF-2K/CaM interactions under non-denaturing conditions (24). Recombinant eEF-2K purified from *E. coli* is fully phosphorylated at T348 (eEF-2K*^p^*); co-expression with λ-phosphatase yields the unphosphorylated form (eEF-2K^λ^). Native gel analysis revealed that apo-CaM_IAE_ binds robustly to S500D-eEF-2K*^p^* (Fig. 1A, 1SA). In contrast, no binding was apparent for eEF-2K^λ^, S500D-eEF-2K^λ^, or eEF-2K*^p^*. To visualize weak apo-CaM_IAE_ binders, we optimized the assay by increasing eEF-2K concentrations and utilizing ADP, which enhances the affinity of eEF-2K for CaM (25). The presence of ADP induced a weak, but observable signal for eEF-2K*^p^* but not for eEF-2K^λ^, suggesting that T348 phosphorylation results in enhanced affinity for apo-CaM_IAE_ (Fig. S1B). Native gel electrophoresis, followed by Western blotting, allowed for the detection of sub-saturating concentrations of Ca^2+^/CaM and also revealed enhanced binding for eEF-2K*^p^*compared to eEF-2K^λ^ (Fig. S1C). We observed a similar enhancement for S500D-eEF-2K*^p^* relative to eEF-2K*^p^*in the presence of Ca^2+^/CaM (Fig S1C). These findings are consistent with previous kinase assays, which have shown that the S500D mutation lowers the apparent *K*_CaM_ (the concentration of CaM required for half-maximal activity) by ∼20 and ∼25-fold in the presence and absence of Ca²⁺, respectively (26). Pre-incubation of S500D-eEF-2K*^p^*with λ-phosphatase before native gel analysis removed *p*T348 as detected by western blot and reduced binding to both apo-CaM and Ca^2+^/CaM (Fig. S1D). Together, our data indicate that T348 and S500 phosphorylation each promote CaM binding, with the maximum effect being observed when both are present.

**Fig. 1.**
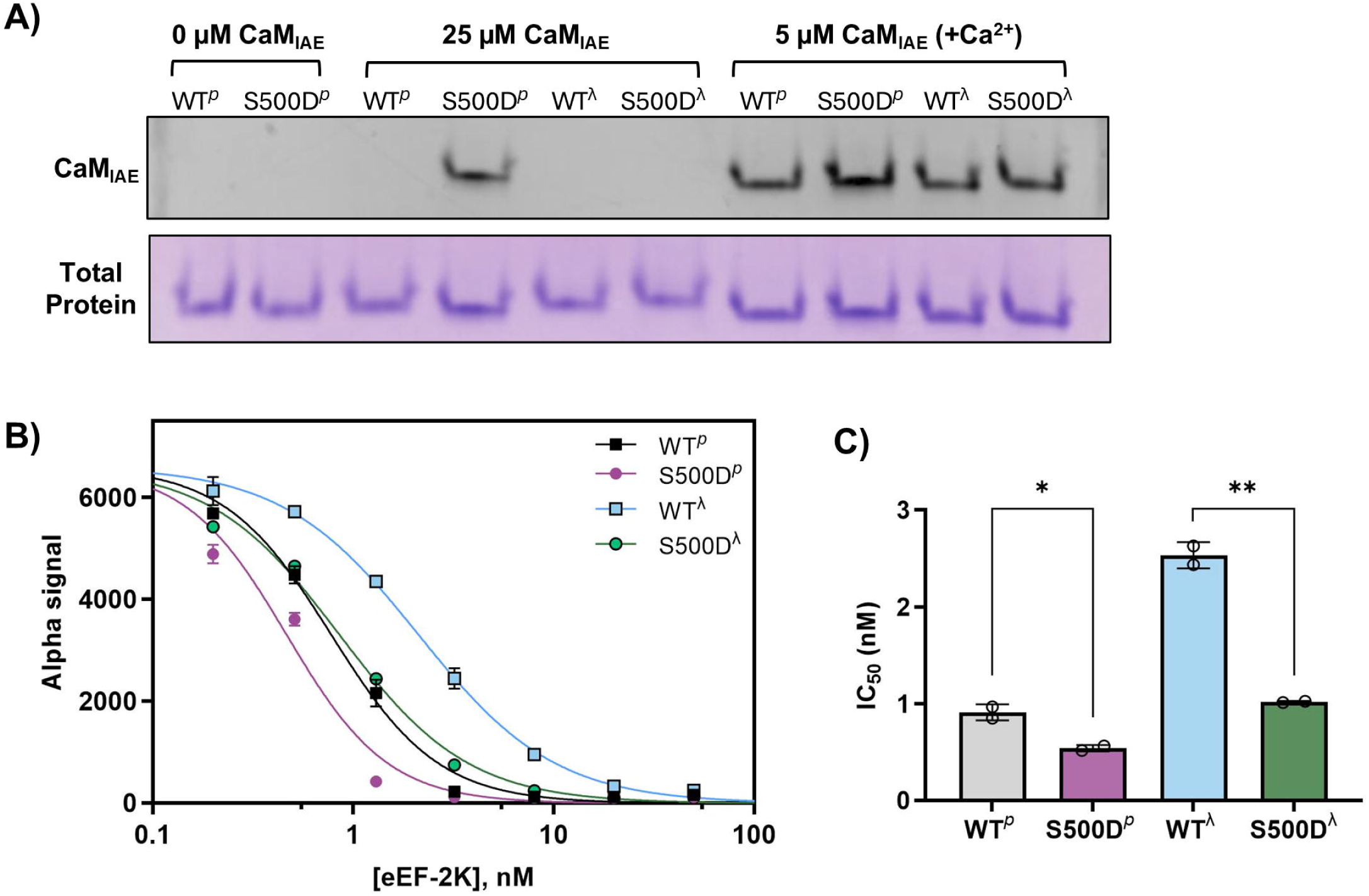
Dual modification of the eEF-2K regulatory loop enhances CaM binding. **(A)** The ability of recombinant wild-type eEF-2K (WT) and the phosphomimetic S500D to bind CaM was assessed via their association with a CaM labeled with the fluorophore IAEDANS (CaM_IAE_) on a nondenaturing gel. The purified proteins were either co-expressed with l-phosphatase (indicated by “^λ^”; unphosphorylated at T348) or not (“*^p^*” indicates phosphorylated on T348). Each construct was incubated with the indicated concentrations of CaM_IAE_. The +Ca^2+^ samples contained 150 μM free Ca^2+^. The samples were run on a native gel to separate the unbound CaM_AE_ from the eEF-2K/CaM_AE_ complexes. The fluorescence of the bound CaM_IAE_ was visualized with a UV imager, and the total protein was determined by Coomassie staining. **(B)** An AlphaScreen assay was used to monitor competition between GST-eEF-2K (10 nM) and untagged eEF-2K mutants (0–50 nM) for His-tagged CaM (2 nM) in the presence of 1 mM Ca^2+^. **(C)** IC_50_ values from the plot in (B) were determined for WT*^p^* (0.78 ± 0.044 nM), S500D*^p^* (0.47 ± 0.026 nM), WT^λ^ (2.1 ± 0.14 nM), and S500D^λ^ (0.86 ± 0.054 nM). Two replicates were used in each case with the error bars representing the standard deviation.

As an additional method to measure CaM binding more quantitatively in the presence of saturating Ca^2+^, we adapted an AlphaScreen-based competition assay (27). GST-tagged eEF-2K constructs were immobilized on anti-GST acceptor beads, while His_6_-tagged CaM was captured on donor beads. Competition with untagged eEF-2K variants enabled the determination of IC_50_ values for CaM binding. Consistent with our native gel assays, phosphorylation of T348 resulted in a ∼2-fold lower IC_50_ for eEF-2K*^p^*compared to the unphosphorylated enzyme (eEF-2K^λ^). The dually modified S500D-eEF-2K*^p^* exhibited a ∼4-fold lower IC_50_ compared to eEF-2K^λ^ (Figs. 1C, D; Table 1), consistent with these two modifications working cooperatively to enhance CaM binding. The IC_50_ values obtained from the AlphaScreen assay for eEF-2K^λ^ and eEF-2K*^p^* are approximately an order of magnitude lower than *K*_d_ values measured previously using solution-based assays for eEF-2K^λ^ and eEF-2K*^p^* (14). This likely reflects the avidity effects inherent to the assay format. Hence, the IC_50_ values reported here may underestimate the magnitude of the effects of these individual or combined modifications. As a negative control, the W85A-eEF2K^λ^ mutant harboring a substitution in the CaM-targeting motif (CTM) (28, 29) displayed a ∼39-fold increase in IC_50_ relative to eEF-2K^λ^ (Fig. S2), validating the sensitivity of the assay to differences in the CaM-binding ability.

**Table 1.**
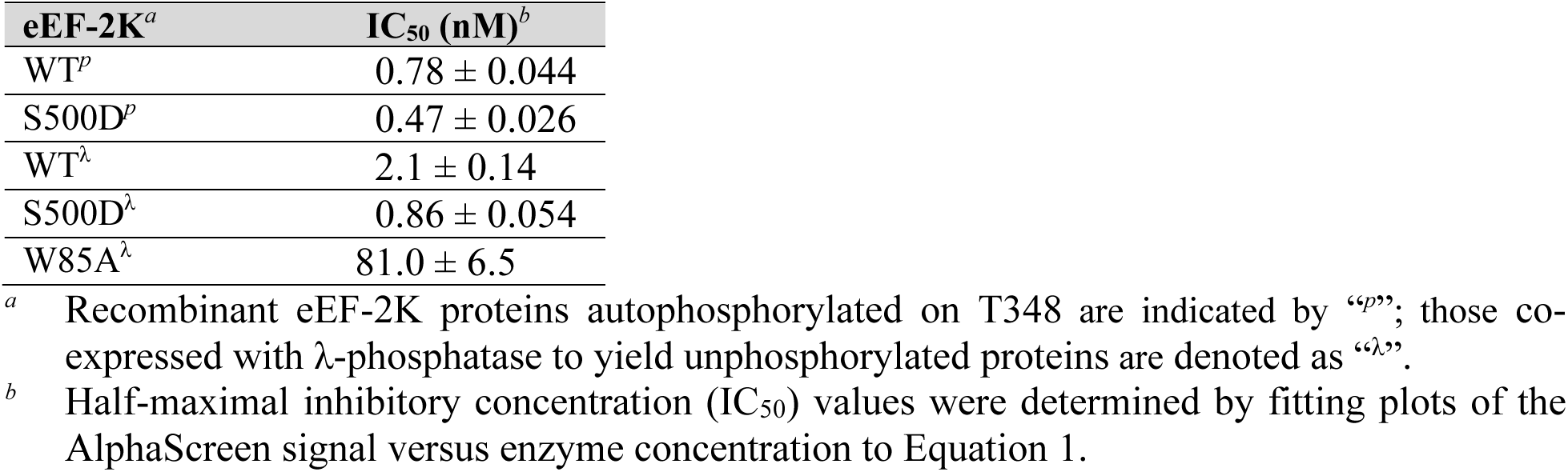
Ca^2+^/CaM binding for recombinant wild-type (WT) and mutant proteins assessed by AlphaScreen.

| **eEF-2K<sup>a</sup>** | **IC<sub>50</sub> (nM)<sup>b</sup>** |
| --- | --- |
| WT<sup>p</sup> | 0.78 ± 0.044 |
| S500D<sup>p</sup> | 0.47 ± 0.026 |
| WT<sup>λ</sup> | 2.1 ± 0.14 |
| S500D<sup>λ</sup> | 0.86 ± 0.054 |
| W85A<sup>λ</sup> | 81.0 ± 6.5 |

<sup>a</sup> Recombinant eEF-2K proteins autophosphorylated on T348 are indicated by “<sup>p</sup>”; those co-expressed with λ-phosphatase to yield unphosphorylated proteins are denoted as “<sup>λ</sup>”.
<sup>b</sup> Half-maximal inhibitory concentration (IC<sub>50</sub>) values were determined by fitting plots of the AlphaScreen signal versus enzyme concentration to Equation 1.

### S500D acts cooperatively with T348 phosphorylation to enhance the intrinsic activity of eEF-2K

We previously reported that the S500D mutation enhances the rate of autophosphorylation at T348 (by ∼6-fold), suggesting that this modification alters the enzyme’s intrinsic properties beyond simply enhancing CaM affinity (26). To assess whether S500 phosphorylation affects the inherent catalytic activity of eEF-2K, we compared the CaM-independent (intrinsic) activities of eEF-2K*^p^* and S500D-eEF-2K*^p^* using a peptide substrate, Pep-S (Fig. 2A, B; Table 2). While the S500D mutant exhibited no significant change in substrate affinity (*K*_m_), it displayed a ∼25-fold increase in intrinsic activity relative to eEF-2K*^p^*, with *k*_cat_ values of 1.1 s^-1^ and 0.04 s^-1^, respectively. Consistent with previous findings, the S500D mutation did not affect the activity (*k_obs_*) of eEF-2K in the presence of saturating Ca²⁺/CaM (Fig. 2C; Table 2).

**Fig. 2.**
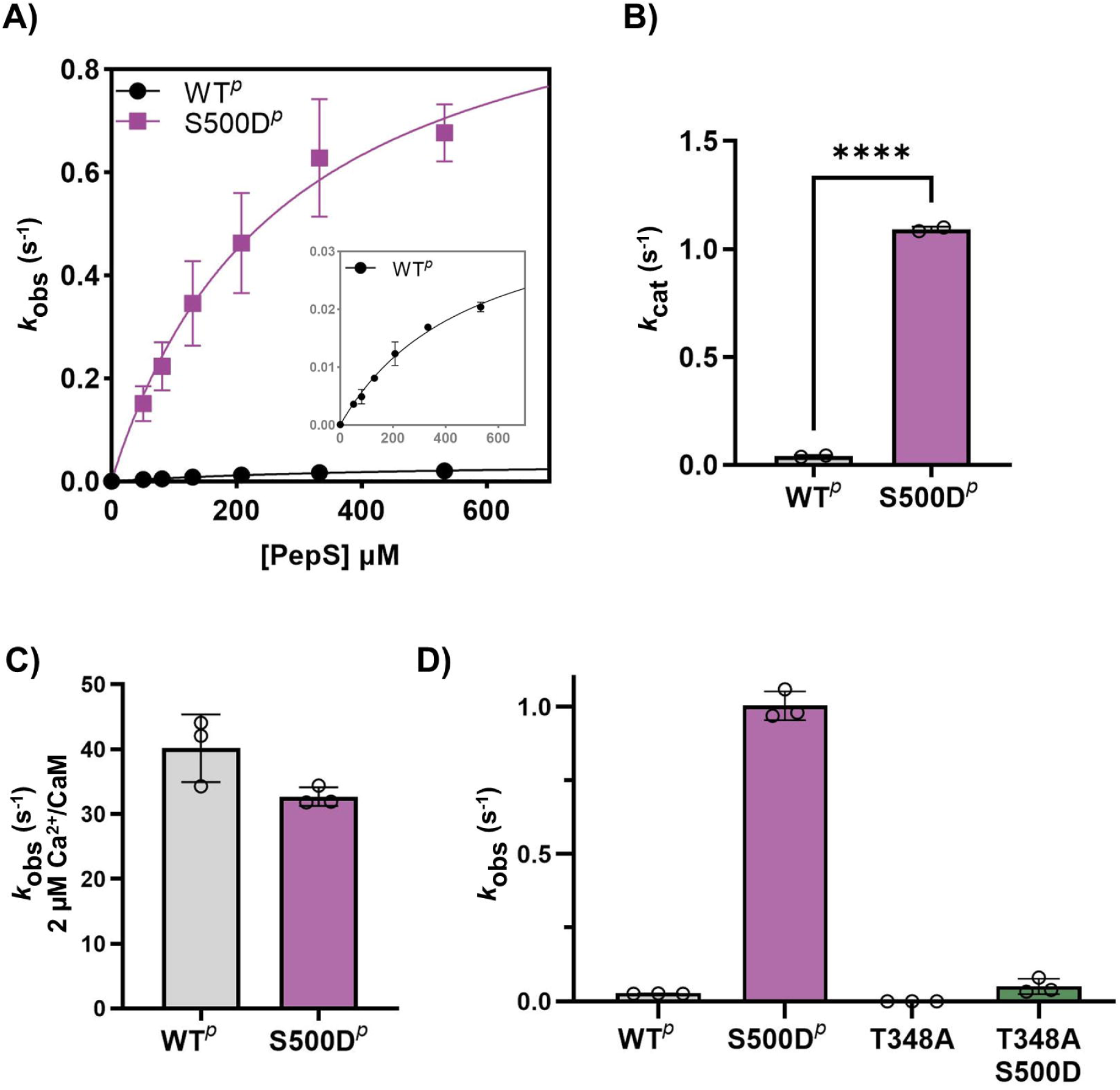
S500D conditionally enhances the catalytic activity of eEF-2K. **(A)** The intrinsic activities of eEF-2K*^p^* and S500D-eEF-2K*^p^* were measured in the absence of CaM using a peptide substrate (Pep-S). The observed rates were plotted versus Pep-S concentration and fitted to the Michaelis-Menten equation. The inset shows a magnification of the y-axis for the eEF-2K*^p^* curve. The estimated *K*_m_ and *k*_cat_ values were 499 ± 86 μM and 0.04 ± 0.004 s^-1^, respectively, for eEF-2K*^p^*and 275 ± 73 μM and 1.0 ± 0.1 s^-1^ for S500D-eEF-2K*^p^*. **(B)** The *k*_cat_ values for duplicate experiments from (A) are displayed as bar graphs. The ‘****’ indicates p < 0.0001 in a t-test. **(C)** The Ca^2+^/CaM-dependent activities of eEF-2K*^p^* and S500D-eEF-2K*^p^* were measured using 450 µM Pep-S to obtain *k*_obs_ values of 40 s^-1^ and 33 s^-1^, respectively. **(D)** The intrinsic activities of eEF-2K*^p^*, S500D-eEF-2K*^p^*, T348A-eEF-2K, and S500D/T348A-eEF-2K were assessed at a single peptide substrate concentration (450 μM Pep-S). The bar graphs display the average of the observed rates (*k*_obs_) in triplicate, which were 0.03 ± 0.005 s^-1^, 1.0 ± 0.06 s^-1^, 0.0007 ± 0.0002 s^-1^, and 0.05 ± 0.03 s^-1^, for eEF-2K*^p^*, S500D-eEF-2K*^p^*, T348A-eEF-2K, and S500D/T348A-eEF-2K, respectively.

**Table 2.** Kinetic parameters for recombinant wild-type (WT) and mutant eEF-2K proteins.

| eEF-2K<sup>a</sup> | Calmodulin<sup>b</sup> | $k_{obs}$<sup>c</sup><br>s<sup>-1</sup> | $k_{cat}$<sup>d</sup><br>s<sup>-1</sup> | $K_m$<sup>d</sup><br>μM |
| --- | --- | --- | --- | --- |
| WT<sup>p</sup> | - | 0.03 ± 0.005<sup>e</sup> | 0.04 ± 0.004 | 499 ± 86 |
|  | + | 40.0 ± 5 | ND<sup>f</sup> | ND |
| S500D<sup>p</sup> | - | 1.0 ± 0.06<sup>e</sup> | 1.0 ± 0.1 | 275 ± 73 |
|  | + | 33.0 ± 2 | ND | ND |
| T348A | - | 0.0007 ± 0.0002 | ND | ND |
| S500D/T348A | - | 0.05 ± 0.03 | ND | ND |
| Δ497-502<sup>p</sup> | - | 0.8 ± 0.1 | ND | ND |

<sup>a</sup> Recombinant eEF-2K proteins contain autophosphorylation at T348, denoted as “<sup>p</sup>”.
<sup>b</sup> Experiments contained 0 or 2 μM calmodulin. All contained ~34 μM free Ca<sup>2+</sup>.
<sup>c</sup> Apparent rate constants ( $k_{obs}$ ) determined in triplicate with 450 μM peptide substrate and 1 mM γ-<sup>32</sup>P-ATP.
<sup>d</sup> Apparent catalytic constants ( $k_{cat}$ ) and apparent Michaelis constants ( $K_m$ ) values were determined from fitting peptide dependence curves to Equation 2.
<sup>e</sup> Average of two sets of triplicates
<sup>f</sup> ND: not determined

To determine whether the S500D mutation can enhance intrinsic activity in the absence of phosphorylation on T348, we utilized the T348A mutant that was previously shown to significantly impair CaM-stimulated activity, reducing both *k*_cat_/*K*_m_ (∼30-fold) towards Pep-S, as well as the corresponding CaM sensitivity (an increase in *K*_CaM_ by ∼7-fold) (14). As expected, in the absence of CaM, a T348A mutation resulted in a ∼20-fold reduction in the rate compared to S500D-eEF-2K*^p^*(Fig. 2D; Table 2). However, the CaM-independent activity of the T348A/S500D double mutant was still ∼70-fold higher than that of T348A (Fig. 2D; Table 2).

These results demonstrate that S500D enhances intrinsic catalytic activity independently of T348 phosphorylation. However, reminiscent of its role in CaM binding, maximal CaM-independent activity is achieved when both T348 and S500 are phosphorylated. Remarkably, the CaM-independent activity of dual-modified enzyme (S500D-eEF-2K*^p^*) is only ∼30-fold less compared to its Ca²⁺/CaM-stimulated form (Table 2).

### CaM binding and S500D may independently disrupt an inhibitory structure around S500

Our previous hydrogen/deuterium exchange mass spectrometry (HXMS) analysis of a truncated eEF-2K construct (eEF-2K_TR_; Fig. S3) in complex with Ca²⁺/CaM revealed reduced deuterium uptake near the CaM-binding site on the N-lobe of the kinase domain and around the phosphate-binding pocket (PBP) that is engaged by T348 upon its phosphorylation (29). Notably, Ca²⁺/CaM binding alone, even in the absence of T348 phosphorylation, also conferred protection in and around the PBP (30), suggesting that CaM binding induces a broader range of conformational changes. To further investigate how CaM binding influences eEF-2K structural dynamics, we performed HXMS measurements on an eEF-2K construct containing an intact R-loop but lacking the 70 N-terminal residues (eEF-2K_ΔN_; unphosphorylated at T348; Fig. S3), in the presence and absence of Ca²⁺/CaM. Among the regions with sufficient peptide coverage, the only statistically significant change in deuterium uptake was observed in a segment spanning residues V502–D513, immediately adjacent to S500. In this region, apo-eEF-2K_ΔN_ exhibited reduced deuterium incorporation compared to its CaM-bound state (Fig. 3A), indicating increased solvent accessibility upon CaM binding, consistent with a conformational rearrangement around S500.

**Fig. 3.**
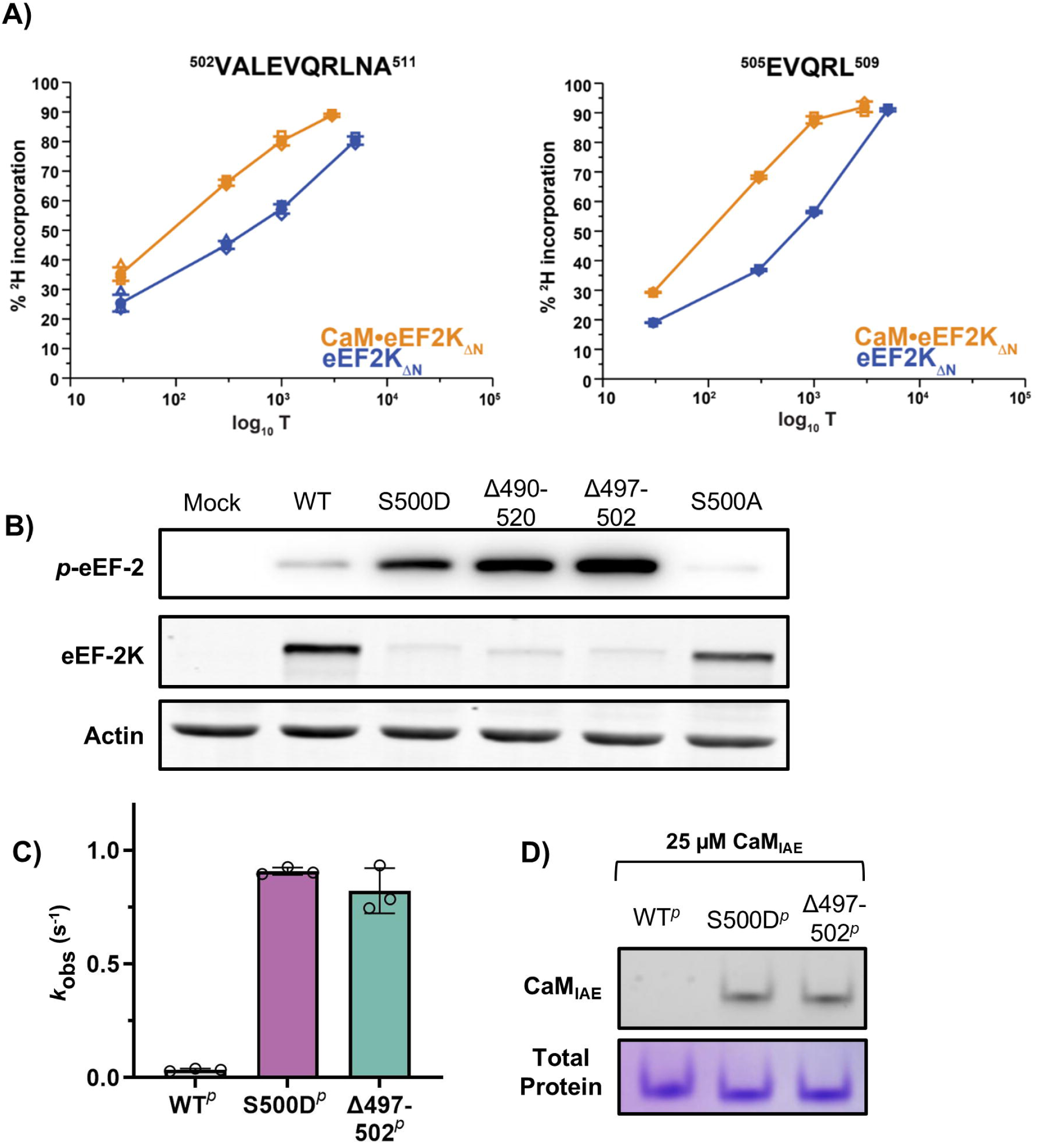
Activation of eEF-2K is associated with disruption of a region around S500. **(A)** Hydrogen-deuterium exchange mass spectrometry (HXMS) experiments were performed on eEF-2K_ΔN_ alone or in a complex with CaM. T348 was dephosphorylated in all cases. The only region within eEF-2K_ΔN_ that showed statistically significant differences between its CaM-free and CaM-bound states involved the V502-D513 segment proximal to S500. Two overlapping peptides corresponding to this region are shown. These peptides illustrate reduced ^2^H incorporation in the CaM-free compared to the CaM-bound state. (**B)** MCF10A eEF-2K^-/-^ were transfected with the indicated eEF-2K mutants. eEF-2K expression and eEF-2 phosphorylation were assessed via western blotting. **(C)** The intrinsic activities of eEF-2K*^p^*, S500D-eEF-2K*^p^*, and eEF-2K_Δ497-502_*^p^* were measured (in duplicate) using 450 μM PepS. *k*_obs_ values of eEF-2K*^p^*, S500D-eEF-2K*^p^*, and eEF-2K_Δ497-502_*^p^* are displayed in bar graph form. The mean values in the three cases were 0.04 s^-1^, 0.9 s^-1^, and 0.8 s^-1^, respectively. **(D)** The association of recombinant eEF-2K*^p^* and corresponding mutants with the fluorescent CaM_IAE_ via native gel.

The above findings led us to hypothesize that the region surrounding S500 contains an inhibitory segment that is displaced upon Ca²⁺/CaM binding or S500 phosphorylation. Biochemical data from the S500D mutant support this model, suggesting that disruption of this region promotes kinase activation. To test this further, we examined the activity of S500D-eEF-2K and two deletion mutants in mammalian cells (Fig. 3B). The expression of S500D-eEF-2K resulted in elevated eEF-2 phosphorylation, despite reduced eEF-2K protein levels, consistent with previous reports suggesting that this mutant is less stable in cells (26, 31). Similarly, a mutant in which the N490–K520 region was replaced by a 6-glycine linker (eEF-2K_Δ490–520_; Fig. S3), showed increased eEF-2 phosphorylation in cells despite reduced protein (Fig. 3B). A more conservative deletion, replacing the P497-V502 region with a 6-glycine linker (eEF-2K_Δ497 502_; Fig. S3), also mimicked the effects of S500D (Fig. 3B).

To further characterize the biochemical properties of the eEF-2K_Δ497-502_ mutant, we expressed and purified it to assess its CaM-independent intrinsic activity against Pep-S *in vitro*. The corresponding T348-phosphorylated form (eEF-2KΔ_497–502_*^p^*) exhibited an intrinsic rate (*k*_obs_ = 0.8 s⁻¹), comparable to S500D-eEF-2K*^p^* (*k*_obs_ = 1.0 s⁻¹) and substantially higher than eEF-2K*^p^* (0.03 s⁻¹) (Fig. 3C; Table 2). Additionally, eEF-2K_Δ497–502_*^p^* displayed a similar capacity to bind apo-CaM as the S500D-eEF-2K*^p^*variant (Fig. 3D). This suggests that this truncation mutant recapitulates key biochemical properties of the dual phosphorylated enzyme.

Collectively, these results support a model in which the disruption of the region surrounding S500 at the C-terminal end of the R-loop, via phosphorylation, mutation, or deletion, relieves an inhibitory constraint, allowing partial activation of eEF-2K and enhancing its ability to bind CaM. Precisely how this is achieved in structural terms remains unclear.

## Discussion

eEF-2K integrates diverse cellular cues, including Ca^2+^ transients and phosphorylation, to regulate translation elongation. Our study reveals that phosphorylation at S500, a site targeted by both PKA (20, 21, 23) and autophosphorylation (15), fine-tunes eEF-2K’s sensitivity to Ca²⁺/CaM by stabilizing a partially active kinase conformation. This modification enhances both the enzyme’s intrinsic (CaM-independent) activity and its affinity for CaM, positioning S500 as a regulatory switch that adjusts kinase activity in response to dynamic cellular conditions.

Our findings suggest that S500 phosphorylation lowers the threshold for activation by Ca²⁺/CaM, enabling eEF-2K to respond to otherwise subthreshold signals. Mechanistically, S500 phosphorylation does not increase eEF-2K activity beyond that when fully saturated with Ca²⁺/CaM (26). Instead, it promotes the formation and stability of the active CaM•eEF-2K complex. This effect is most pronounced when S500 phosphorylation is combined with T348 autophosphorylation. The doubly modified enzyme (S500D/T348-phosphorylated) exhibits a ∼1400-fold increase in intrinsic activity (Table 2) and a ∼100-fold enhancement in the CaM sensitivity (reduced *K*_CaM_) compared to the T348A enzyme (14, 26). Together, these observations suggest that these modifications cooperatively stabilize an active conformation.

HXMS studies reveal that Ca²⁺/CaM binding increases solvent accessibility in the region adjacent to S500, suggesting a conformational rearrangement. Deletion of S500 and nearby residues mimics the effects of a S500D mutation, resulting in robust CaM-independent activity and elevated eEF-2 phosphorylation despite reduced eEF-2K protein levels. These findings support a model in which the S500-adjacent region acts as an inhibitory element that can be relieved either by Ca²⁺/CaM binding or by S500 phosphorylation. We previously demonstrated that T348 phosphorylation binds to a basic pocket in the kinase domain (14, 29), likely inducing a conformational change within the R-loop. Dual phosphorylation at T348 and S500 may stabilize an open, CaM-bound-like conformation that lowers the energetic barrier for CaM binding, enhancing eEF-2K’s affinity for both apo- and Ca^2+^-bound CaM. Importantly, S500 phosphorylation also accelerates the rate of T348 autophosphorylation by ∼6-fold (26), further reinforcing its role as a priming modification that promotes eEF-2K activation and stabilizes the active complex. Thus, phosphorylation at these two sites, T348 and S500, situated at opposite ends of the R-loop, synergize to stabilize an active state.

Recent studies using a CaM-eEF-2K fusion protein show that covalent CaM attachment renders kinase activity insensitive to Ca²⁺ (24), indicating that Ca²⁺ primarily enhances CaM binding. However, S500 autophosphorylation remains strictly Ca^2+^-dependent (26), suggesting that Ca²⁺ induces specific conformational changes required to enable this modification. In wild-type eEF-2K, efficient S500 autophosphorylation requires both Ca²⁺/CaM binding and prior T348 phosphorylation and proceeds with much slower kinetics than T348 phosphorylation (∼10 minutes versus ∼0.3 seconds) (26). This suggests that S500 autophosphorylation functions as a delayed regulatory step, integrating sustained Ca²⁺/CaM signaling. In contrast, PKA-mediated phosphorylation of S500 bypasses the Ca²⁺/CaM requirement, providing a mechanism for global signals (e.g., nutrient status, energy stress, or cAMP pathway activation) to modulate eEF-2K sensitivity (23, 26). PKA-mediated S500 phosphorylation strengthens CaM affinity, reducing the Ca²⁺/CaM threshold for activation and heightening eEF2K responsiveness under basal conditions. Notably, the *K*_CaM_ for S500D-eEF-2K*^p^* (∼1.5 μM) (26) for apo-CaM falls within the physiological range of cellular CaM concentrations (2–25 μM) (32), suggesting that the doubly phosphorylated enzyme may be receptive to activation by CaM that is mostly Ca²^+^-free at basal Ca²⁺ levels (∼100 nM) (33). In addition to these phosphorylation-dependent effects, our previous work has demonstrated that ADP binds to eEF-2K and promotes its interaction with CaM (25). This suggests that ADP may act in conjunction with S500 phosphorylation as an allosteric modulator, enhancing the assembly of the active CaM•eEF-2K complex, particularly under conditions of metabolic stress or low ATP availability.

Phosphorylation at S500 has also been implicated in promoting the ubiquitin-mediated degradation of eEF-2K, the cause of the reduced protein levels in the S500D mutant (see, for example, Fig. 3B) (31). This highlights a dual function for the modification: while it enhances kinase activity and facilitates CaM binding, it may simultaneously serve as a degradation signal, marking eEF-2K for proteasomal turnover. Importantly, S500 phosphorylation alone is not sufficient to promote degradation. Abrogation of CaM binding or kinase activity rescues S500D eEF-2K expression in cells, suggesting that degradation of S500D is only induced in the context of the active eEF-2K•CaM complex (26). Such a mechanism could provide tight temporal control over eEF-2K activity by coupling activation with self-limiting inactivation, thereby preventing sustained signaling. This type of dual-purpose regulation, where a single post-translational modification both activates and earmarks a protein for degradation, contrasts with the regulatory paradigm observed in CaMKII (34–36). Although CaMKII phosphorylation at T286 increases CaM affinity similarly and enables partial Ca²⁺-independent activity, it is not known to promote degradation. In contrast, S500 phosphorylation in eEF-2K appears to promote activation and degradation pathways, revealing a dynamic, self-regulating control of the kinase in response to synaptic activity, metabolic stress, and growth factor signaling.

Collectively, our findings support a model (illustrated schematically in Fig. 4) in which S500 phosphorylation, T348 autophosphorylation, and Ca²⁺/CaM binding converge to stabilize the active conformation of eEF-2K. Through this mechanism, eEF-2K modulates translation elongation in response to diverse cellular inputs, thereby fine-tuning the proteome to match the cell’s metabolic and signaling demands.

**Fig. 4.**
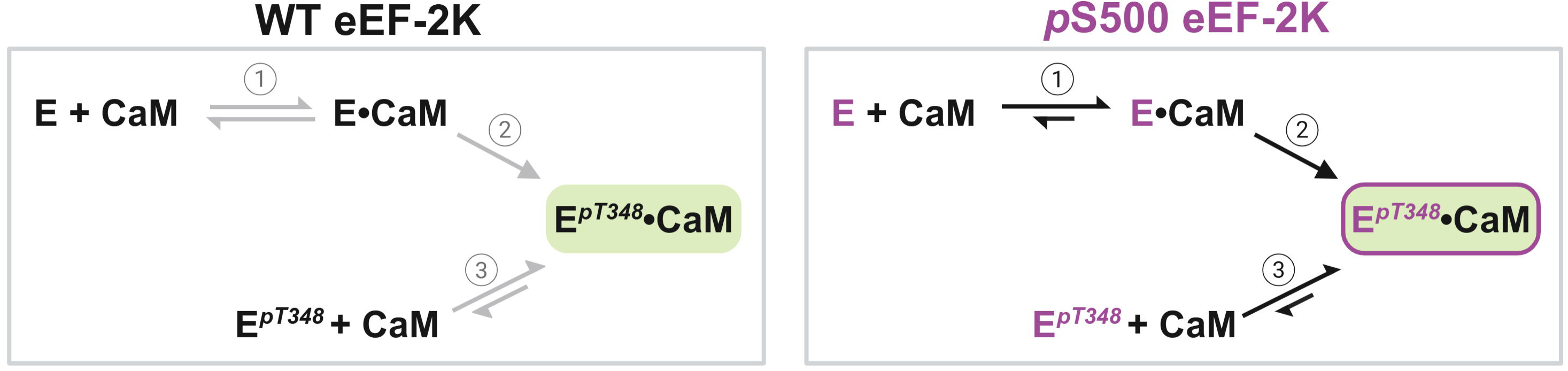
Proposed mechanism for the regulation of eEF-2K by S500 phosphorylation. *Left panel:* Wild-type eEF-2K (E, black) has low activity until bound to CaM (E•CaM). The formation of this complex (step 1) promotes autophosphorylation at T348 (step 2) to yield the active complex (E*^p^*^T348^•CaM). The E*^p^*^T348^ form has a higher affinity for CaM than E and partially stabilizes the active complex (step 3). *Right panel:* Phosphorylation of S500 (mimicked by the S500D mutant, E, purple) increases the amount of active complex (E*^p^*^T348^•CaM) by increasing the affinity of E for CaM (step 1), enhancing the rate of T348 autophosphorylation (step 2) and increasing the affinity of the corresponding E*^p^*^T348^ form for CaM (step 3). Cellular eEF-2K is phosphorylated at S500 either through CaM-induced autophosphorylation or by PKA. In the former case, phosphorylation on S500 occurs once T348 is phosphorylated and serves to trap eEF-2K in the E*^p^*^T348^•CaM complex by significantly enhancing CaM affinity, i.e., affecting only step 3. PKA-mediated phosphorylation of S500, however, can influence all three steps to drive the formation of a stable E*^p^*^T348^•CaM complex.

Further studies are needed to define the structural basis of these regulatory mechanisms more completely. The absence of a full-length eEF-2K structure limits our understanding of how the presence of an intact R-loop modifies the interactions of T348 and S500 with the kinase core, and possibly with CaM. Furthermore, the physiological relevance of S500 phosphorylation in specific contexts, such as neurons where eEF-2K is crucial for synaptic plasticity (9, 11, 37), remains to be elucidated. Our findings provide a mechanistic framework for exploring how post-translational modifications, CaM dynamics, and signaling pathways converge to regulate eEF-2K function in health and disease.

### Experimental Procedures

#### Purification of recombinant proteins

Recombinant eEF-2K and corresponding mutant proteins were expressed and purified using a protocol similar to that described previously (38). Briefly, the eEF-2K vector (pET32a Trx-His_6_-eEF-2K) was grown in Rosetta-gami 2 (DE3) competent cells (Novagen) at 37 °C until an optical density (OD) of 0.6-0.8 was reached. The cells were then induced with 0.5 mM IPTG and collected after 16 h of expression at 22 °C. To purify eEF-2K without phosphorylation on T348, eEF-2K was co-expressed with λ-phosphatase (pCDF-Duet1). The Trx-His_6_-tagged proteins were purified from the cell lysate using Ni-NTA affinity chromatography, and the tags were removed using TEV protease. The proteins were further purified using an ÄKTA pure FPLC system equipped with a Mono Q 10/100 GL anion exchange column (Cytiva) and a HiPrep 26/60 Sephacryl S-200 HR gel filtration column (Cytiva). Purified eEF-2K was dialyzed into storage buffer (25 mM HEPES, pH 7.5, 50 mM KCl, 0.1 mM EDTA, 0.1 mM EGTA, 2 mM DTT, and 10% glycerol), concentrated, and stored at −80°C. CaM and IAEDANS-labeled CaM were purified as described previously (39).

#### Native gel-based assays

The ability of recombinant eEF-2K proteins (0.67 μM) to bind 25 μM apo-CaM_IAE_ or 5 μM Ca^2+^/CaM (150 μM free Ca^2+^) was assessed by incubating samples for 20 min at room temperature in Native Binding Buffer (25 mM HEPES (pH 6.8), 50 mM KCl, 1 mM EGTA, 2 mM DTT, 0.005% Brij, and native sample buffer (Bio-Rad #1610738)) and 0 or 1.15 mM CaCl_2_ in a final volume of 30 µL. The samples were loaded onto a nondenaturing gel (Bio-Rad Mini-PROTEAN TGX, 4-15%) in Native Running Buffer (25 mM Tris, 192 mM glycine) and run at 50-80V, 4 °C, protected from light. The CaM_IAE_ fluorescence was measured using a GE Amersham Imager 600 (excitation at 312 nm). The total protein was visualized by Coomassie staining.

To monitor the binding of recombinant eEF-2K proteins with subsaturating concentrations of Ca^2+^/CaM, native gels were analyzed by western blotting. Samples contained 0.2 µM eEF-2K and 0.2 µM CaM, with 150 µM free calcium in Native Binding Buffer (see above), at a final volume of 40 µL. The samples were allowed to incubate for 20 minutes at room temperature before being loaded onto a nondenaturing gel, as described above. The native proteins were transferred to a PVDF membrane for 16 h at 30V in 25 mM Tris, 192 mM glycine, 2 mM CaCl_2_, and 20% methanol and then probed with CaM (4830, Cells Signaling Technology), eEF-2K (C-12, sc-390710, Santa Cruz Biotechnology), or *p*T348 eEF-2K (EP4411, ECM Biosciences) specific antibodies.

#### AlphaScreen assay

Anti-His_6_ Alpha donor beads (#AS116M) and AlphaLisa anti-GST acceptor beads (#AL110M) were purchased from Revvity. The assay buffer consisted of 25 mM HEPES (pH 7.5), 50 mM KCl, 10 µM EGTA, 10 µM CaCl_2_, 2 mM DTT, 0.02% (w/v) Tween-20, and 0.01% (w/v) BSA. All samples were processed in white 384-well plates (Corning, #3825). 10 nM GST-tagged eEF-2K (SignalChem, #E01-10G-320) and 2 nM His_6_-tagged CaM (EMD Millipore, #208670) with 1 mM excess CaCl_2_ were incubated with 0 – 50 nM of eEF-2K^λ^, S500D-eEF-2K^λ^, eEF-2K*^p^* or S500D-eEF-2K*^p^*, or 0 – 5 µM W85A-eEF-2K^λ^ for 30 min at room temperature. A mixture of donor and acceptor beads (5 ng/µL) was added to each sample, mixed by centrifugation at 800 rpm for 3 min, and incubated for 2 h at room temperature, protected from light. The alpha signal was detected using a BioTek Synergy Neo2 multi-mode microplate reader (Agilent). All concentrations above are given as final concentrations in 10 µL total sample volumes.

Data were fitted to Equation 1 to obtain IC_50_ values, where “max” represents the maximum AlphaScreen signal and “min” is the signal after complete inhibition. The maximum and minimum values were set to be equal for all datasets, and the Hill slopes (n) were allowed to vary.

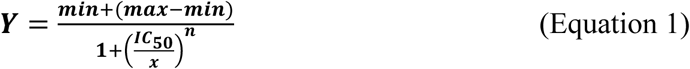

#### Kinase assays

CaM-independent intrinsic activity assays were performed by combining 400 nM eEF-2K*^p^*, 40 nM S500D-eEF-2K*^p^*, 800 nM T348A-eEF-2K, or 400 nM T348A/S500D-eEF-2K with either a single concentration, 450 mM, of Pep-S (Ac-RKKYKFNEDTERRRFL-Amide, Peptide 2.0), or varied concentrations of Pep-S in Activity Buffer containing 25 mM HEPES (pH 7.5), 2 mM DTT, 20 μg/mL BSA, 100 μM EDTA, 100 μM EGTA, 10 mM MgCl_2_, 0.005% Brij, 50 mM KCl, and 150 µM CaCl_2_. Reactions were initiated with 1 mM [γ-^32^P]-ATP, and time points were taken and analyzed as described previously (15). CaM-dependent assays at a single CaM concentration were carried out with 1 nM eEF-2K*^p^*or S500D-eEF-2K*^p^*, 150 mM PepS, and 2 μM CaM in Activity Buffer. For single peptide concentrations, data were fit by linear regression. For Pep-S dependence curves, the data were fit to Equation 2.

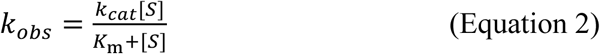

#### Hydrogen-deuterium exchange mass spectrometry measurements

CaM and eEF-2K_ΔN_ were expressed and purified using protocols described previously (40). eEF-2K_ΔN_ was treated twice with a ∼1/100 ratio of λ-phosphatase overnight at 4 °C to obtain fully dephosphorylated species. The CaM-bound eEF-2K_ΔN_ sample was prepared by incubating eEF-2K_ΔN_ in a buffer containing 20 mM Tris (pH 7.5), 200 mM NaCl, 2 mM DTT, 1.0 mM CaCl_2_ with a threefold excess of CaM in an identical buffer and then purifying the respective complexes by size exclusion chromatography, as previously described (40).

Stock solutions (∼45 μM) were diluted ∼35-fold in an otherwise identical D_2_O-based buffer (pH* = 7.5) and incubated for variable amounts of time at 15 °C. The exchange was quenched by diluting the sample 1/1 with a solution containing 5 M guanidinium chloride and 1% formic acid at 2 °C. The sample was then digested using an Enzymate BEH Pepsin column (Waters). The resulting peptides were captured using a C8 Trap Cartridge and then resolved using a C18 HPLC column, inline with a maXis-II ETD ESI-QqTOF spectrometer (Bruker). More details on the experimental protocol have been published elsewhere (40). Reference spectra were obtained by diluting the samples in a H_2_O-based buffer and then collecting MS/MS data, which were subsequently analyzed using Bruker COMPASS and BIOTOOLS software packages. HDExaminer 3.3.0 (Sierra Analytics) was used to assess ^2^H-incorporation. Two datasets were collected on different dates using similar but distinct samples. In the first data collection, a single experiment was recorded for the following time points - eEF-2K_ΔN_: 30 s, 300 s, 1000 s, and 5000 s; CaM/eEF-2K_ΔN_ complex: 30, 300s, 1000 s, and 3000 s. In the second data collection, two time points (30 s, 300 s) were collected in triplicate for all samples. The overall coverage for CaM and eEF-2K_ΔN_ were 88.5% and 64.5%, respectively.

#### Mutant eEF-2K activity in MCF10A cells

Wild-type and mutant eEF-2K (cloned into plasmid 10792; Addgene) were transfected into MCF10A eEF-2K^-/-^ cells as previously described (14). Twenty-four hours after transfection, cells were lysed, and protein concentrations were determined using a Pierce™ BCA assay (Catalog number: T23227; Thermo Fisher Scientific). 30 μg of protein was then subjected to SDS-PAGE and western blotting as described previously (14). For western blotting, the following antibodies were used: eEF-2K (C-12) (Catalog number: sc-390710;Santa Cruz Biosciences); eEF-2 (Catalog number: 2332; Cell Signaling); phospho-eEF-2 (Catalog number: 2331; Cell Signaling); Pan-actin (Catalog number: 8456; Cell Signaling); IRDye® 800CW Goat anti-Rabbit (Catalog number: 926-32211; LiCor); IRDye® 680RD Goat anti-Mouse (Catalog number: 926-68070; LiCor); and Goat Anti-Rabbit IgG (H+L)-HRP Conjugate (Cat # 1721019, Bio-Rad).

## Acknowledgments

This work is supported by NIH award R01 GM123252 (to KND and RG), the Robert A. Welch Foundation (F-2185, KND), and a CPRIT award RP210088 (to KND).

## Author contributions

A.L.B and L.S.B. designed research, performed biochemical experiments, analyzed data, and generated the manuscript and figures, which were refined by K.N.D and R.G. The AlphaScreen assay was designed by R.M.S and E.J.C., and carried out by R.M.S. HXMS studies were done by A.P. Recombinant proteins for biochemical studies were prepared by A.L.B. and C.D.J.T. and cell biology was performed by L.S.B.

## Conflict of interest

The authors declare that they have no conflicts of interest with the contents of this article.

## This article contains supporting information

**Fig. S1.**
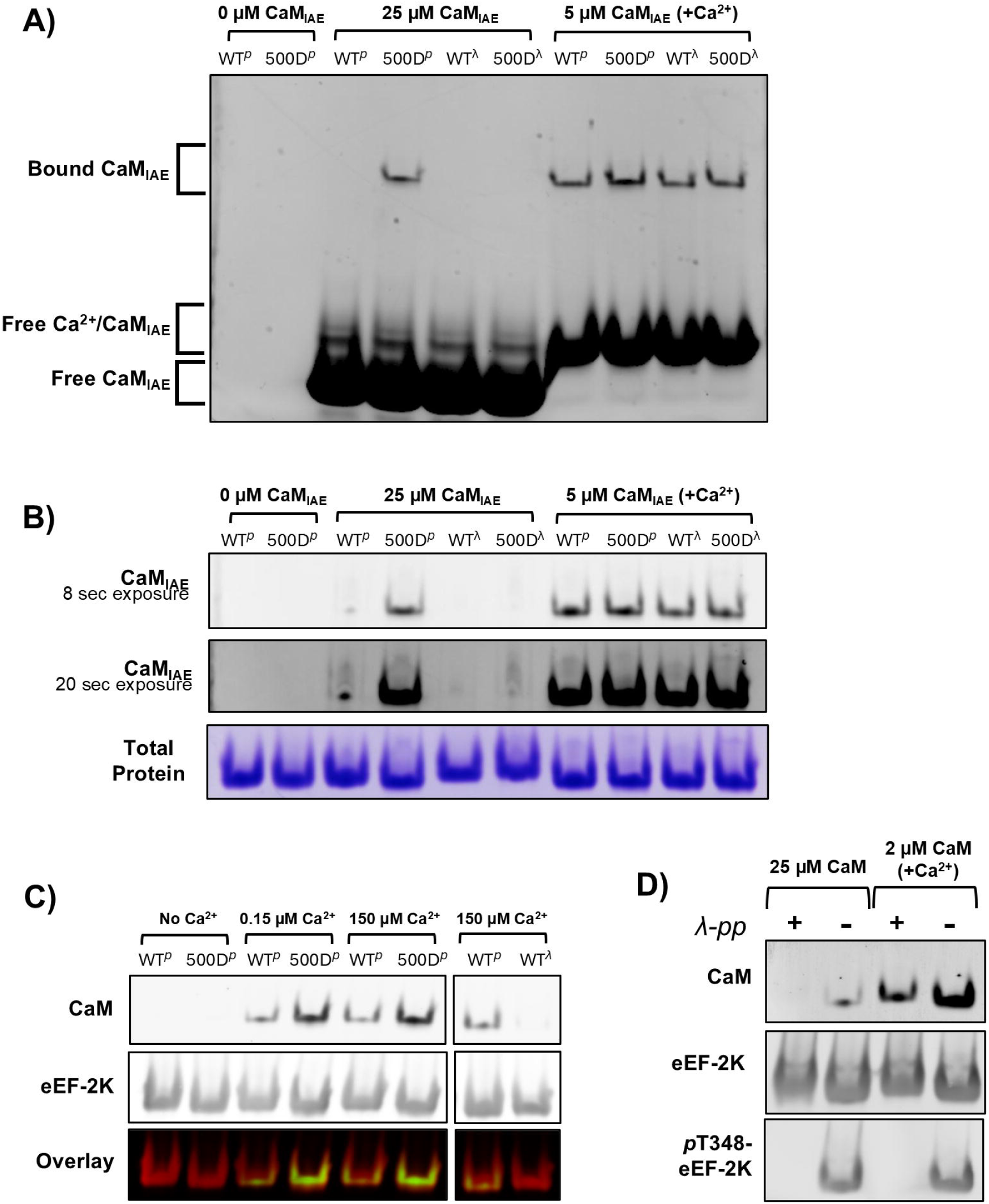
Native gel binding assay with CaM_IAE_. **(A)** The entire gel from Fig. 1B, showing bound and unbound CaM_IAE_ fluorescence (512 nm excitation). The bound CaM_IAE_ comigrates with recombinant eEF-2K (∼82 KDa) on the native gel, while the excess free CaM (∼17 KDa) migrates more rapidly through the gel. Addition of Ca^2+^ causes a slight upward shift in free CaM. **(B)** The ability of recombinant eEF-2K proteins (“*^p^*” denotes presence of *p*T348, while “^λ^” denotes coexpression with λ-phosphatase, i.e., T348 is not phosphorylated) to bind CaM_IAE_ on a native gel was assessed similarly to Fig. 1B, except the incubation tubes contained 1 mM ADP and the enzyme concentration was increased from 0.67 to 2.5 µM. The CaM_IAE_ fluorescence was measured using two different exposure times, 8 and 20 sec, to visualize low fluorescence signals. **(C)** Assessment of binding of recombinant eEF-2K proteins (0.2 µM) to CaM (0.2 µM) by native gel followed by western blotting with antibodies for CaM and eEF-2K. The concentrations of free Ca^2+^ are indicated. **(D)** S500D-eEF-2K*^p^* was pre-incubated with or without purified λ-phosphatase (λ-pp) before combining with either 25 µM calcium-free CaM or 2 µM CaM with 150 µM free Ca^2+^. Samples were loaded onto a native gel and analyzed by Western blot using CaM, eEF-2K, and *pT348-eEF-2K-specific* antibodies.

**Fig. S2.**
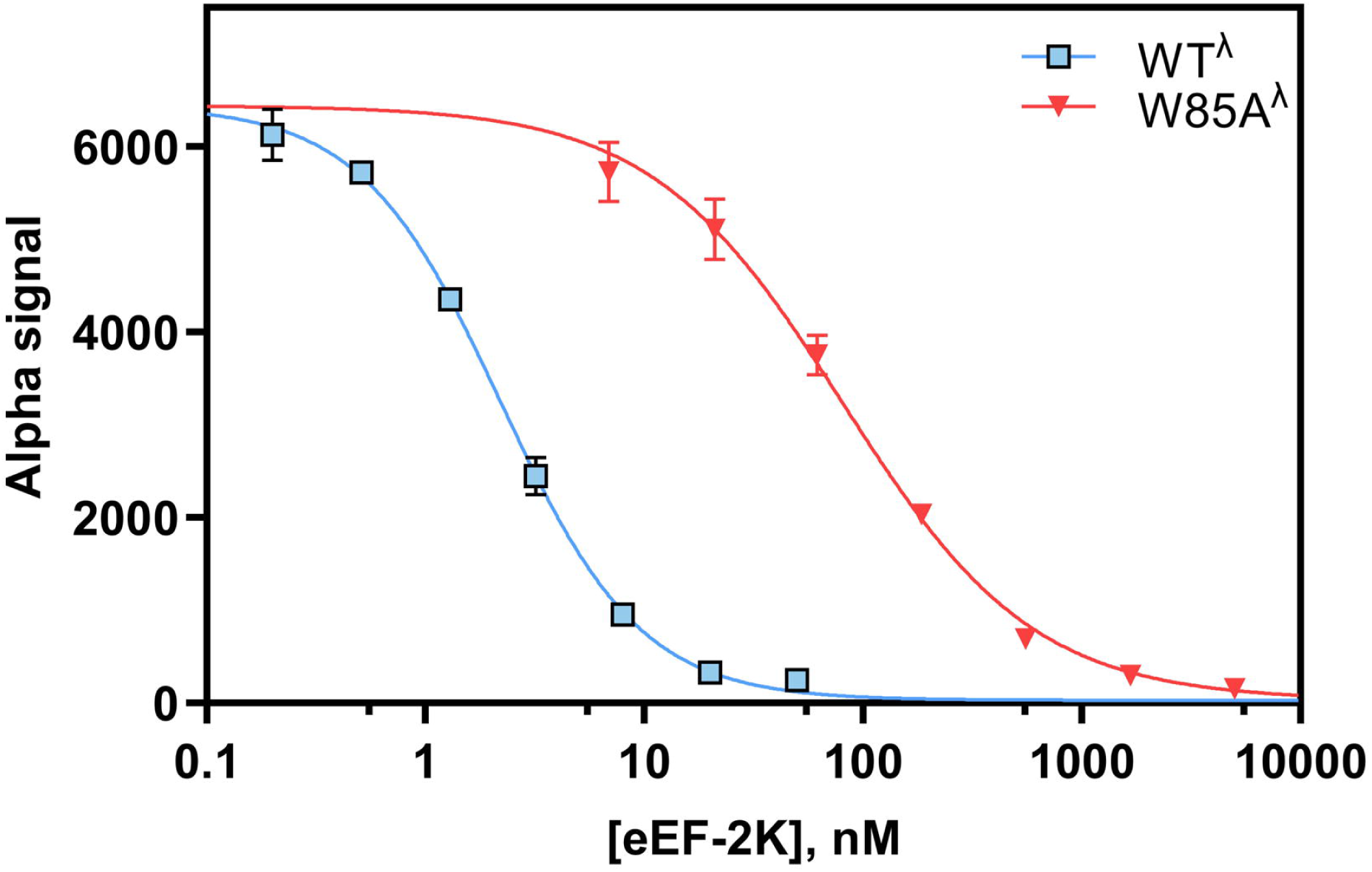
AlphaScreen CaM binding assay for WT^λ^ vs W85A^λ^ eEF-2K. An AlphaScreen assay was used to monitor competition between GST-eEF2K (10 nM) and untagged WT^λ^ and W85A^λ^ eEF2K mutants (0-5 µM) for His_6_-tagged CaM (2 nM) in the presence of 1 μM Ca^2+^. IC_50_ values for WT^λ^ and W85A^λ^ eEF2K were determined to be 2.1 ± 0.14 nM and 81 ± 6.5 nM, respectively. Data were measured in duplicate in all cases; error bars represent standard deviation.

**Fig. S3.**
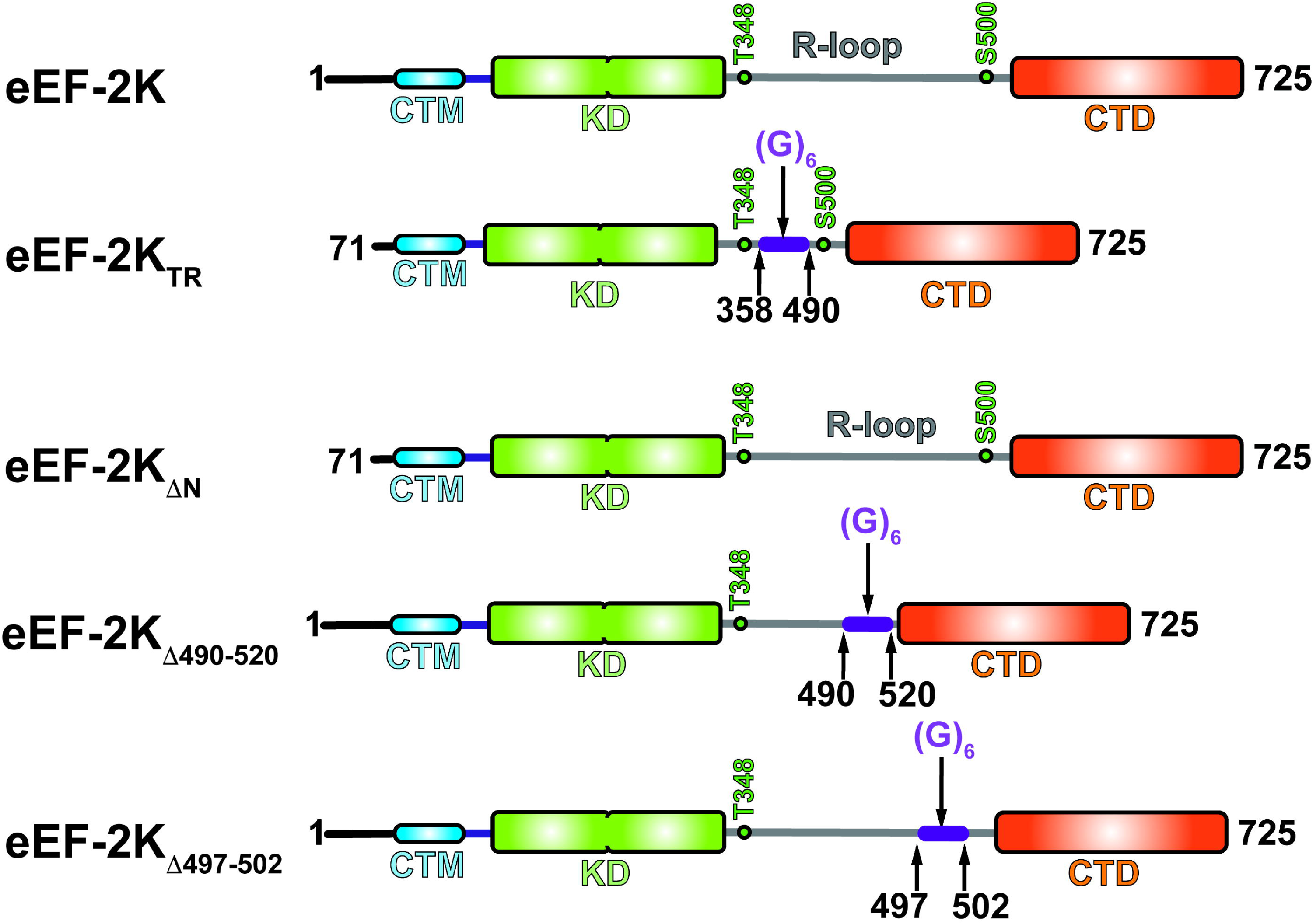
eEF-2K constructs used this study. Schematic representation of the various eEF-2K constructs, including wild-type full-length eEF-2K, the eEF-2K_TR_ (a 6-glycine linker has replaced missing 70 N-terminal residues, and the 359-490 segment; the linker is indicated in purple), eEF-2K_ΔN_ (missing 70 N-terminal residues but containing an intact regulatory loop), eEF-2K_Δ490-520_ (a 6-glycine linker has replaced 490-520 segment), and eEF-2K_Δ497-502_ (a 6-glycine linker has replaced 490-502 segment). The N-terminal calmodulin-targeting motif (CTM), the α−kinase domain (KD), the regulatory loop, and the C-terminal domain (CTD) are indicated. The activating T348 and S500 sites are located at the two ends of the regulatory loop.

## Notes

### Competing Interest Statement

The authors have declared no competing interest.

